# Structural insights into the recognition and catalysis of tRNA by human NSUN2

**DOI:** 10.64898/2026.01.09.698742

**Authors:** Qian Hu, Wen Yang, Yunyun Yu, Ran Yi, Yutong Zhang, Leyuan Duan, Lei Wang, Fudong Li, Kaiming Zhang, Qingguo Gong, Shanshan Li

**Author notes:** Corresponding authors. (K.Z.); (Q.G.); (S.L.).

## Abstract

The human RNA m^5^C methyltransferase NSUN2 catalyzes site-specific cytosine methylation across diverse RNA substrates and thereby regulates a wide range of biological and physiological processes. However, the molecular basis by which NSUN2 achieves broad substrate recognition while maintaining catalytic specificity has remained unclear. Here we determine structures of human NSUN2 in both substrate-free and substrate-bound states using X-ray crystallography and cryo-electron microscopy. Structures of NSUN2 in complex with multiple tRNA substrates reveal a structure-first, sequence-tolerant strategy in which NSUN2 actively remodels tRNA architecture, exposing the buried target cytosine and positioning it within the catalytic pocket for methyl transfer. This recognition strategy enables NSUN2 to accommodate diverse tRNA substrates through a largely conserved interaction interface. Guided by these structural insights, we identify a small-molecule inhibitor that suppresses NSUN2 activity and cancer cell proliferation. Together, our findings define the molecular principles underlying NSUN2-mediated RNA m^5^C modification and provide a structural foundation for targeting NSUN2 in cancers.

## Introduction

5-Methylcytosine (m^5^C) is a widespread post-transcriptional RNA modification found in messenger RNA (mRNA), transfer RNA (tRNA) and ribosomal RNA (rRNA)^1–3^. By regulating RNA stability, nuclear export and translation efficiency, m^5^C plays essential roles in embryonic development, neurogenesis, and cellular stress responses^4^. Dysregulation of RNA m^5^C modification and its regulatory enzymes has been increasingly linked to human diseases, particularly cancers, highlighting the need to understand the underlying molecular mechanisms^5,6^.

In eukaryotes, RNA m^5^C is catalyzed primarily by the NOL1/NOP2/SUN (NSUN) family (NSUN1-NSUN7) and by DNA methyltransferase 2 (DNMT2)^7^. These enzymes are S-adenosylmethionine (SAM)-dependent cytosine methyltransferases that catalyze methyl transfer through a covalent enzyme-RNA intermediate. In NSUN enzymes, catalysis relies on two conserved cysteines that coordinate intermediate formation and methylated product release^8^. Although substrate repertoires and modification sites have been mapped for most NSUN enzymes, mechanistic understanding of substrate recognition remains strikingly limited at the structural level. To date, NSUN6 is the only family member for which substrate-bound structures and detailed recognition mechanisms have been elucidated^9,10^. Structural information for other NSUN proteins is largely restricted to cofactor-bound or protein-protein complexes, providing limited insight into how RNA substrates are engaged and remodelled during catalysis^11,12^.

As a prominent member of the NSUN family, NSUN2 displays an unusually broad substrate spectrum and catalyzes site-specific m^5^C modifications in diverse classes of RNA. These include positions 48-50 at the junction between the variable loop and T-arm of mature tRNAs, position 34 in the anticodon loop of pre-tRNAs, as well as multiple sites within the 5′ untranslated region (UTR), coding sequence (CDS), and 3′ UTR of mRNAs, in addition to other non-coding RNAs^13–16^. Genome-wide sequencing studies have shown that NSUN2-dependent m^5^C sites are highly enriched in the CNGGG motif, which resembles the C48-C50 region of tRNA, suggesting that this G-rich environment may serve as a common anchoring feature for NSUN2 binding^17^. Through these activities, NSUN2 regulates diverse biological processes, including cell differentiation, proliferation and senescence, and its dysregulation has been implicated in breast, bladder, gastric and lung cancers, as well as leukaemia^18–21^. Accordingly, NSUN2 has emerged as both a key regulatory node in RNA metabolism and a potential therapeutic target.

Despite its biological and clinical importance, the structural basis by which NSUN2 recognizes and modifies such a wide range of RNA substrates remains unknown. Here, we address this gap by determining structures of human NSUN2 in both substrate-free and substrate-bound states. Using X-ray crystallography and single-particle cryo-electron microscopy (cryo-EM), we solved the structure of NSUN2 bound to SAH and captured NSUN2 in complex with multiple representative tRNA substrates, including tRNA^Tyr^, tRNA^Lys^ and pre-tRNA^Leu^. These structures reveal a conserved and dynamic mechanism by which NSUN2 recognizes tRNA substrates through shared architectural features and actively induces large-scale RNA conformational rearrangements to expose otherwise buried target cytosines. Furthermore, guided by the structural insights, we performed structure-based screening and identified a small-molecule inhibitor that selectively suppresses NSUN2 activity by disrupting substrate RNA binding. Together, our findings elucidate the molecular basis of NSUN2-mediated tRNA methylation and provide a structural framework for targeting NSUN2 in diseases.

## Results

### Capturing pre- and post-substrate tRNA recognition states of NSUN2

To elucidate the molecular mechanism by which NSUN2 recognizes and modifies its RNA substrates at atomic resolution, we first determined the 2.5-Å resolution crystal structure of full-length human NSUN2 in complex with S-adenosyl-L-homocysteine (SAH), a product analog of the methyl donor S-adenosyl-methionine (SAM) (Fig. 1a,b). NSUN2 adopts a compact, disc-like architecture composed of an N-terminal methyltransferase (MTase) domain and a C-terminal domain assembly. The MTase domain displays a canonical Rossmann fold, with SAH clearly resolved in the catalytic pocket (Fig. 1b). The observed cofactor-binding mode closely resembles that of other SAM-dependent methyltransferases^9^, supporting the functional integrity of the crystallized enzyme. The C-terminal region shows weak structural similarity to PUA RNA-binding domains and forms an extended positively charged surface, suggesting a role in RNA substrate engagement (Extended Data Fig. 1a). We therefore refer to this region as a previously uncharacterized PUA-like RNA-binding module of NSUN2.

**Fig. 1:**
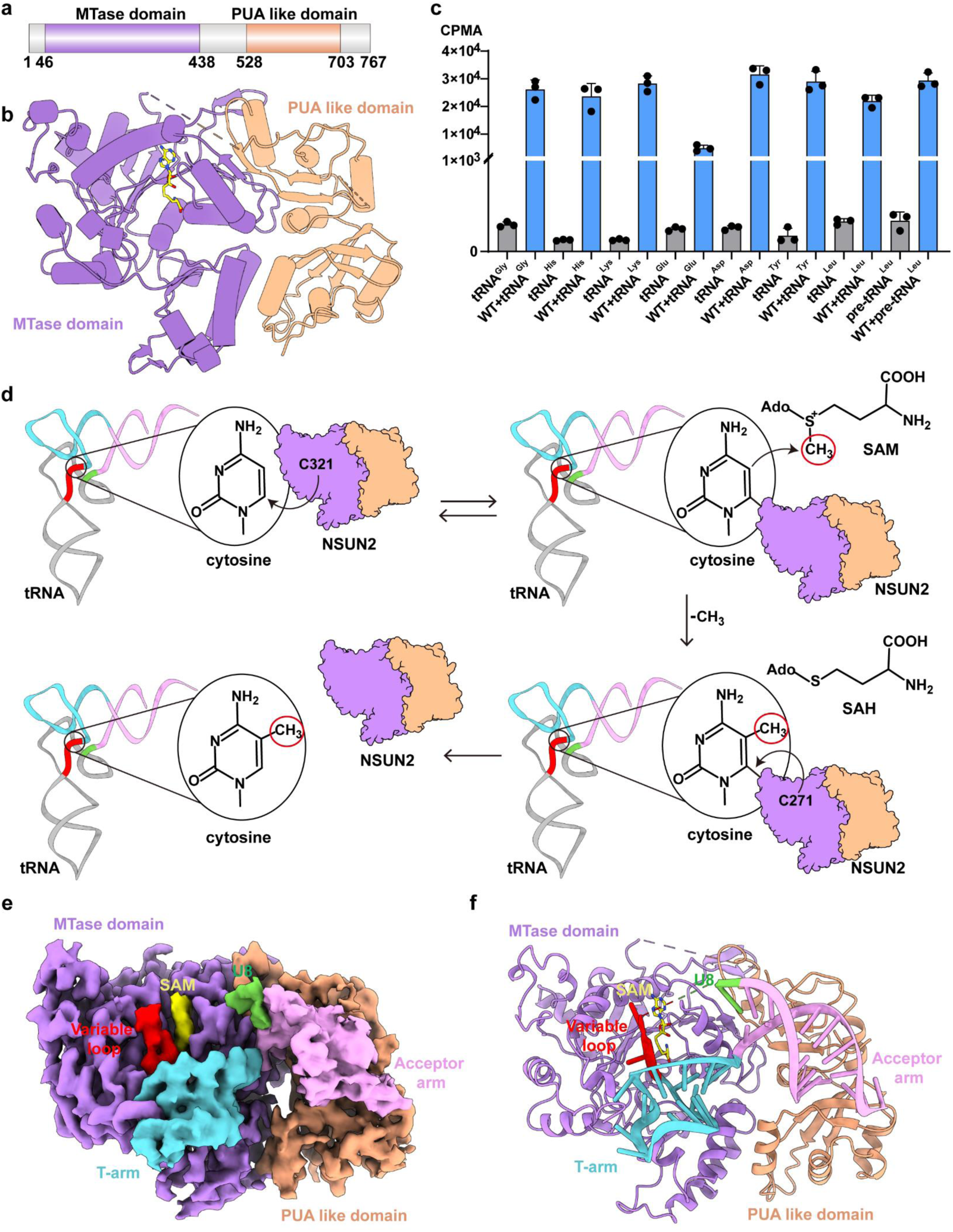
Structural snapshots of NSUN2 before and after tRNA engagement. **a.** Domain organization of NSUN2. MTase domain is shown in purple, the PUA-like domain in orange, and the disordered regions in gray. **b.** The crystal structure of NSUN2 in complex with SAH. **c.** Enzymatic activities of purified NSUN2 with different tRNA substrates. Data are shown as the mean ± SD (n = 3 independent experiments). **d.** Schematic illustration of the NSUN2 catalytic mechanism. **e.** The cryo-EM map of the NSUN2-tRNA^Tyr^-SAM. **f.** The atomic model of the NSUN2-tRNA^Tyr^-SAM.

To assess substrate specificity, we tested NSUN2 activity against seven mature tRNAs and one precursor tRNA *in vitro*. NSUN2 exhibited measurable methyltransferase activity toward all tested tRNAs, with varying efficiencies (Fig. 1c). Among these substrates, tRNA^Tyr^ showed relatively strong activity and was selected for structural analysis. As reported, NSUN2 contains two conserved catalytic cysteines, Cys321 and Cys271, which are predicted to act sequentially during m^5^C formation. Cys321 initiates nucleophilic attack on the C6 position of cytosine to form a covalent enzyme-RNA intermediate that activates the C5 position for methyl transfer from SAM, whereas Cys271 promotes bond cleavage to release the methylated RNA (Fig. 1d). To trap a substrate-bound state, we generated a C271A mutant and reconstituted the NSUN2(C271A)-tRNA^Tyr^-SAM complex. Biochemical assays showed that this mutant forms a stable, SAM-dependent covalent complex with tRNA but lacks catalytic activity, consistent with a post-recognition & pre-methylation intermediate (Extended Data Fig. 1b,c). We next determined its structure by cryo-EM at an overall resolution of 3.34 Å (Fig. 1e,f; Extended Data Fig. 2). The cryo-EM map resolves both the MTase and PUA-like domains, as well as the bound tRNA and cofactor.

Comparison of the substrate-free NSUN2 structure with the substrate-bound cryo-EM structure reveals pronounced conformational changes upon tRNA binding. In particular, helices spanning residues 133-137 and 143-155 shift toward the catalytic center, resulting in a more compact substrate-binding pocket (Extended Data Fig. 1d). Several active-site residues also undergo side-chain rearrangements that position them closer to the target cytosine and SAM (Extended Data Fig. 1e). Together, these structures capture distinct pre- and post-recognition states of NSUN2, providing direct structural evidence for how NSUN2 undergoes coordinated domain and active-site rearrangements to enable tRNA methylation.

### Conformational dynamics of tRNA^Tyr^ during NSUN2 recognition

In its canonical tertiary structure, tRNA adopts an L-shaped conformation stabilized by long-range interactions at the elbow and inner-corner regions^22^. In tRNA^Tyr^, the NSUN2 target cytosine C48 is located within the variable loop and forms a non-canonical base pair with G15 in the D-loop, contributing to stabilization of the inner-corner region of the L-shaped tRNA (Fig. 2a). Access of NSUN2 to C48 therefore requires local disruption of this tertiary interaction network. Consistent with this requirement, the substrate-bound NSUN2(C271A)-tRNA^Tyr^-SAM structure reveals extensive conformational rearrangements of the tRNA upon enzyme binding. The acceptor arm, T-arm and variable loop engage a positively charged surface on NSUN2 and undergo coordinated movements that reposition C48 into the catalytic pocket (Fig. 2b,c). Relative to the predicted free tRNA^Tyr^ structure, the T-arm rotates outward and the acceptor arm tilts downward by ∼20°, increasing the angle between the two arms to ∼117° and disrupting their planar stacking (Fig. 2d,e). These conformational changes disrupt long-range tertiary interactions at the tRNA elbow (Fig. 2f) and inner corner (Fig. 2g). As a result, G46, U47 and C48 undergo base flipping, exposing C48 for entry into the NSUN2 active site. Consistent with conformational flexibility, the D-arm and anticodon arm do not form stable contacts with NSUN2 and are poorly resolved in the cryo-EM density (Extended Data Fig. 3a). To test whether a canonical L-shaped tRNA is required for NSUN2 recognition, we disrupted elbow interactions by introducing G18A, G19A and G18A/G19A mutations in tRNA^Tyr^. Neither binding affinity nor methylation activity was significantly affected by these mutations (Fig. 2h; Extended Data Fig. 3b), indicating that NSUN2 may not require a pre-formed L-shaped tRNA for initial recognition. Together, these results show that NSUN2 binding induces substantial remodeling of tRNA^Tyr^ to expose the target cytosine for methylation.

**Fig. 2:**
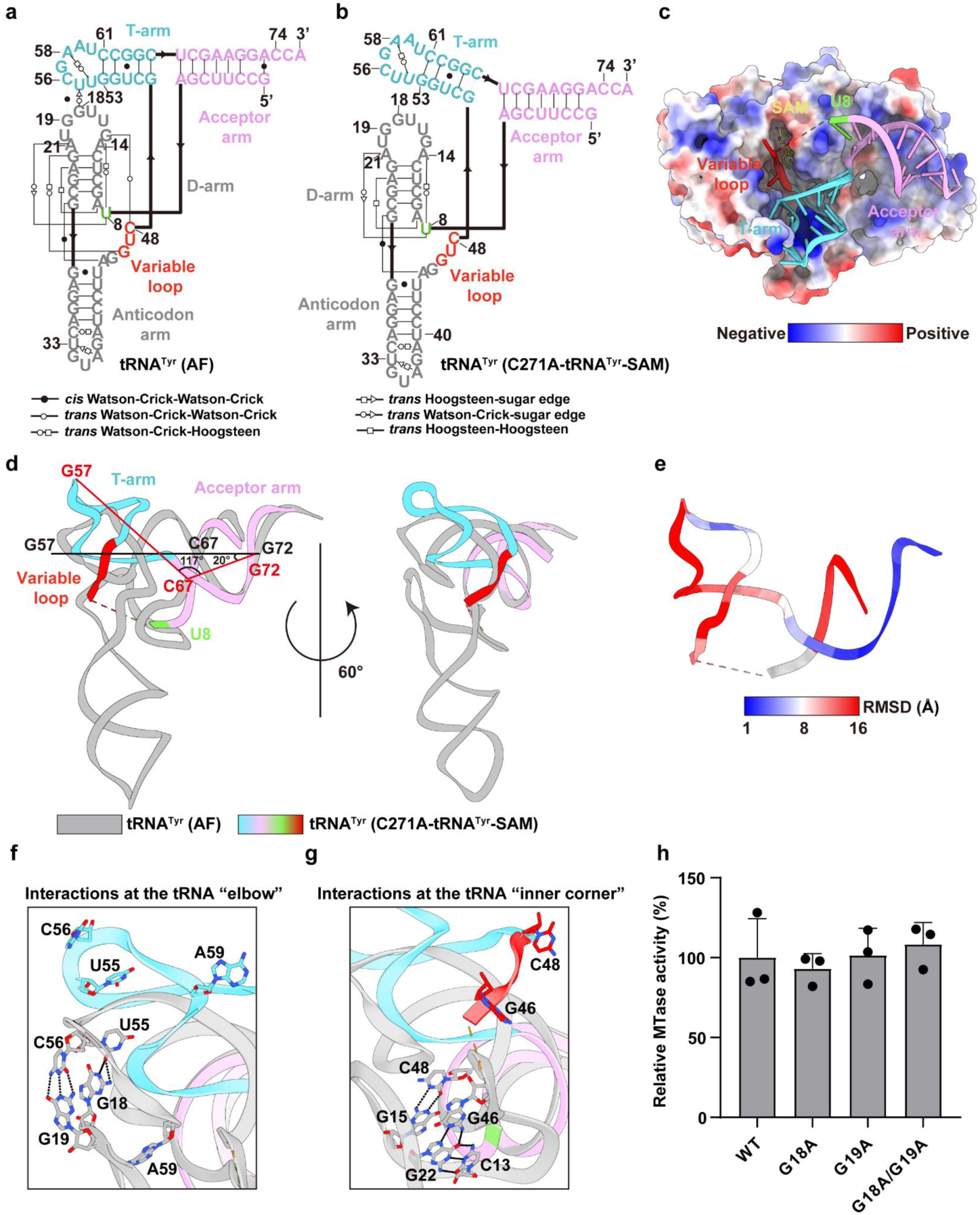
Conformational changes in tRNA^Tyr^ during recognition. a-b. Secondary structure schematic of tRNA^Tyr^ predicted by AlphaFold (a) and that observed in the NSUN2-tRNA^Tyr^-SAM complex (b)**. c.** Binding between the NSUN2 and tRNA^Tyr^. tRNA^Tyr^ is depicted as sticks and NSUN2 with its electrostatic surface. **d-e.** Comparison between the tRNA^Tyr^ predicted by AlphaFold and observed in the NSUN2-tRNA^Tyr^-SAM complex (d). The tRNA^Tyr^ in the NSUN2-tRNA^Tyr^-SAM complex is colored by C-alpha RMSD in ChimeraX (e). **f-a. g.** Detailed-view interactions of the elbow and inner corner in tRNA^Tyr^ predicted by AlphaFold and observed in the NSUN2-tRNA^Tyr^-SAM complex. **h.** Enzymatic activities of purified NSUN2 with various tRNA mutants. Data are shown as the mean ± SD (n = 3 independent experiments).

### Molecular interfaces mediating NSUN2-tRNA^Tyr^ binding

In the substrate-bound state, NSUN2 engages tRNA^Tyr^ through multiple interfaces distributed across the MTase domain and the PUA-like domain (Fig. 3a, b). At the global level, the MTase domain contacts the acceptor arm-D-arm junction, the T-arm and the variable loop, whereas the PUA-like domain interacts exclusively with the acceptor arm (Fig. 3a). This organization suggests distinct roles in substrate recognition and catalysis.

**Fig. 3:**
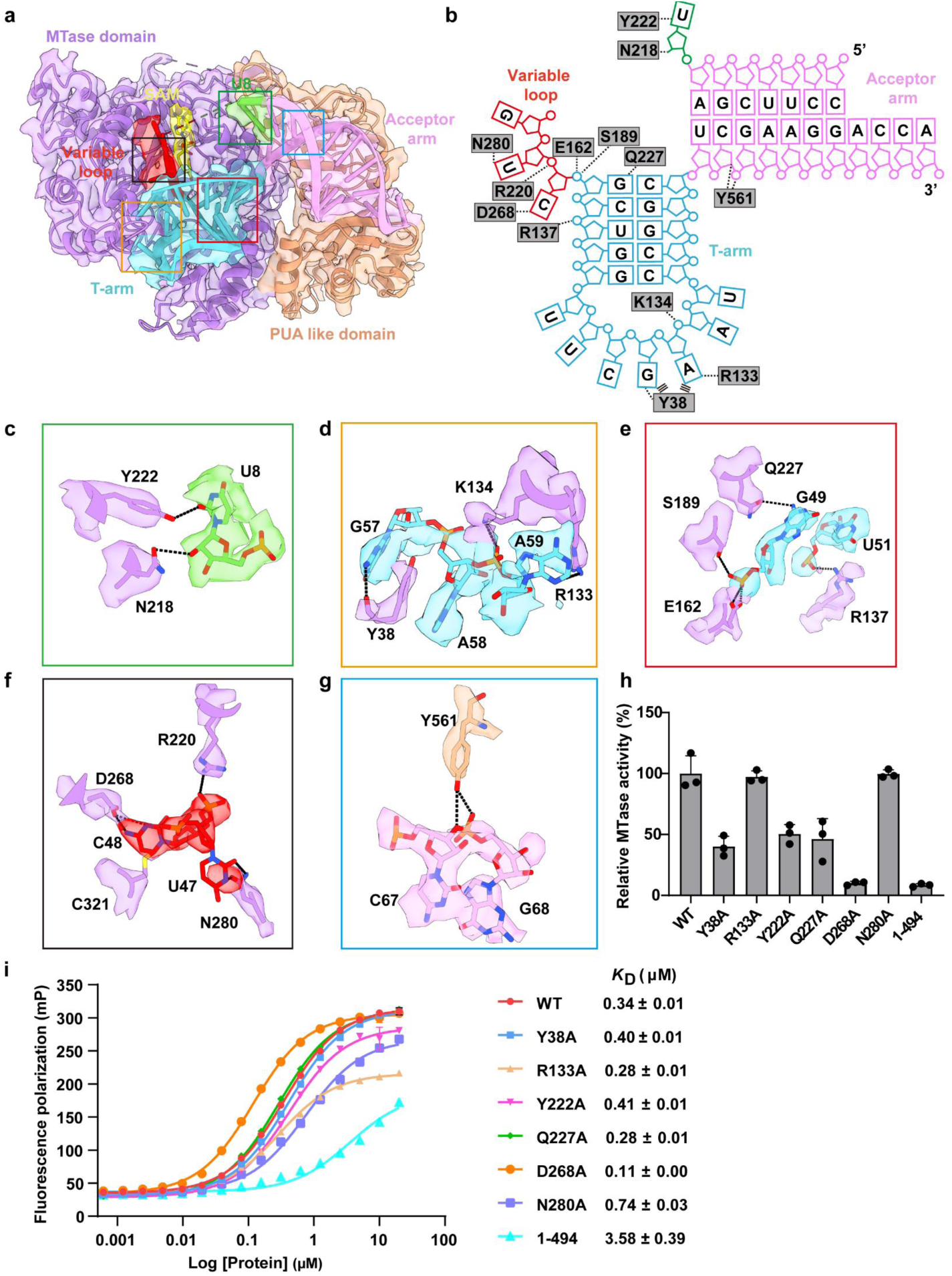
Binding interfaces between NSUN2 and tRNA^Tyr^. **a.** The cryo-EM map and atomic model of the NSUN2-tRNA^Tyr^-SAH complex, illustrating the interaction interfaces. **b.** Schematic representation of NSUN2 interactions with the tRNA^Tyr^. Hydrogen bonds are shown with black dashed lines, and π-stacking interactions with black lines. **c-g**. Detailed view of NSUN2-tRNA^Tyr^ interactions. **h**. Comparison of relative activities of wild-type (WT) and variant NSUN2 methylating tRNA^Tyr^. Data are shown as the mean ± SD (n = 3 independent experiments). **i.** Comparison of tRNA^Tyr^ binding affinities between WT and mutant NSUN2 measured by fluorescence polarization. Data are shown as the mean ± SD (n = 3 independent experiments).

Within the MTase domain, SAM occupies the catalytic pocket in a configuration similar to that observed in the NSUN2-SAH crystal structure (Extended Data Figs. 1d, 1e, 4a). The adenine ring is accommodated in a hydrophobic pocket, while surrounding polar residues coordinate the ribose and terminal groups (Extended Data Fig. 4a). Alanine substitutions of these residues markedly reduced methyltransferase activity, confirming their functional importance (Extended Data Fig. 4b). Adjacent to the catalytic pocket, the MTase domain establishes interactions with the acceptor arm-D-arm junction. Y222 forms a base-specific interaction with U8, and N218 contacts the U8 ribose (Fig. 3b, c). Consistent with a role in substrate catalysis rather than initial binding, the Y222A mutation strongly reduced catalytic activity with minimal effect on binding affinity (Fig. 3h, i). The MTase domain further stabilizes the T-arm through a network of base- and backbone-mediated interactions. R133 and Q227 contact A59 and G49, respectively, while S189 and E162 engage the phosphate backbone of G49 (Fig. 3b, d, e). Mutational analysis showed that Q227 is required for efficient catalysis, while R133 primarily contributes to substrate binding (Fig. 3h, i). Notably, tRNA binding induces ordering of the NSUN2 N-terminal region. Residues 36-41, which are disordered in the substrate-free NSUN2 structure, become structured in the tRNA-bound complex and participate directly in T-arm recognition (Extended Data Fig. 1d). In particular, Y38 inserts between G57 and A58 via π-π stacking, and its backbone contacts G57 (Fig. 3d). The Y38A mutation retained binding but reduced activity to ∼40% of wild-type (Fig. 3h, i). At the catalytic center, the variable loop inserts directly into the MTase pocket. G46, U47 and the target cytosine C48 are resolved in the active site, where N280 contacts U47 and D268 forms base-specific interactions with C48 (Fig. 3b, f). Mutational analysis showed that D268 is essential for catalysis, whereas N280 mainly contributes to substrate binding (Fig. 3h, i). Finally, the contribution of the PUA-like domain was assessed using a truncated NSUN2 construct containing only the MTase domain (residues 1-494). This truncation strongly reduced both binding affinity and catalytic activity (Fig. 3h, i). Structural analysis indicates that this domain anchors the acceptor arm through phosphate-mediated interactions, including contacts between Y561 and the ribose and phosphate groups of C67 and G68 (Fig. 3g). Together, these interactions define a coordinated binding mode in which the MTase domain mediates base-specific recognition and catalysis, while the PUA-like domain stabilizes tRNA engagement.

**Fig. 4:**
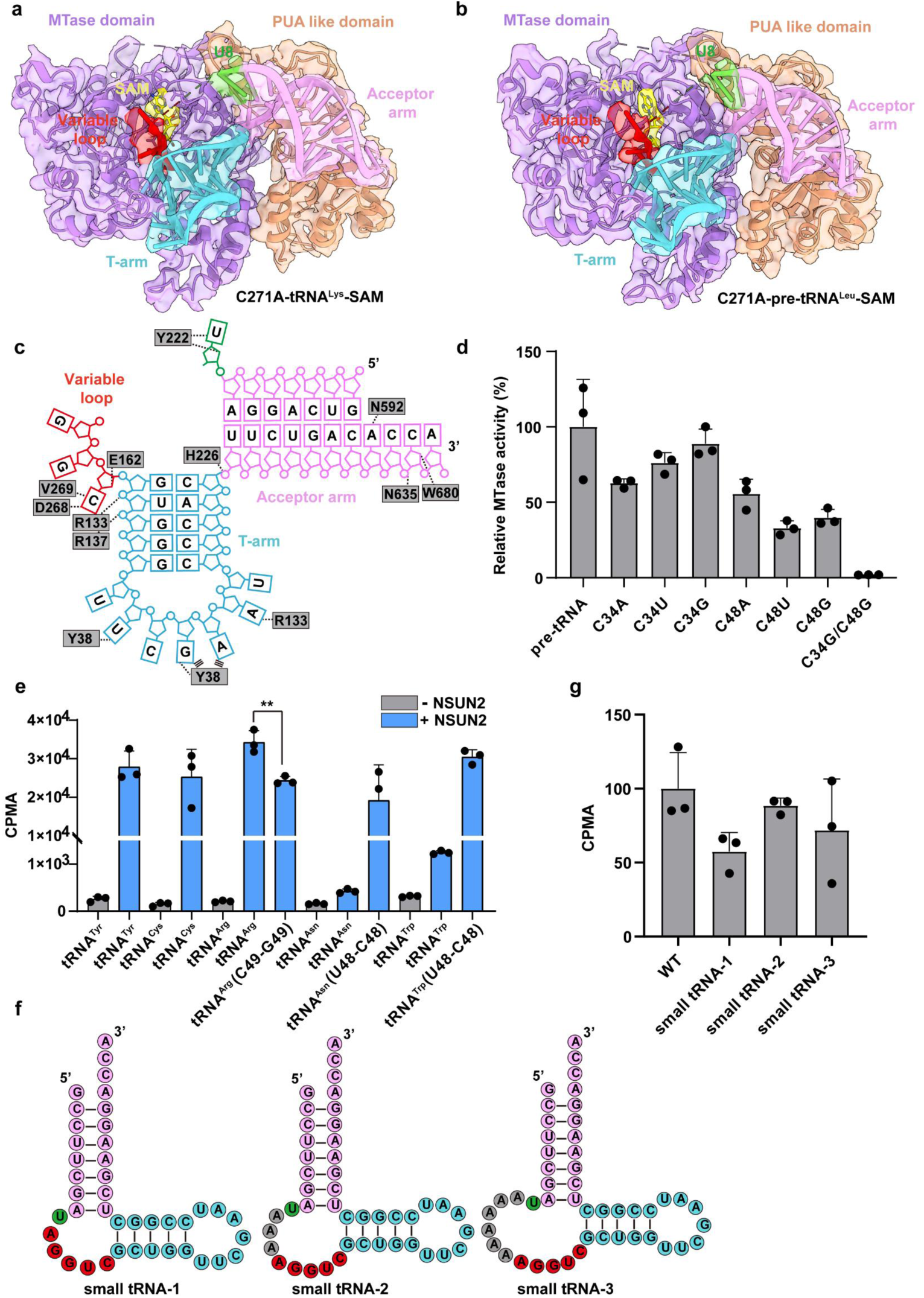
Conserved Mode of Recognition for a Broad Range of tRNA Substrates by NSUN2. a-b. The cryo-EM map and atomic model of the NSUN2-tRNA^Lys^-SAM complex (a) or the NSUN2-pre-tRNA^Leu^-SAM complex (b). **c.** Schematic representation of NSUN2 interactions with the pre-tRNA^Leu^. Hydrogen bonds are shown with black dashed lines, and π-stacking interactions with black lines. **d.** Comparison of relative activities of WT and variant pre-tRNA^Leu^ methylated by NSUN2. Data are shown as the mean ± SD (n = 3 independent experiments). **e.** Enzymatic activities of purified NSUN2 with different tRNA substrates or various tRNA mutants. Data are shown as the mean ± SD (n = 3 independent experiments). **f.** Secondary structure diagrams of three small tRNAs. **g.** Enzymatic activities of purified NSUN2 with various small tRNAs. Data are shown as the mean ± SD (n = 3 independent experiments).

### A conserved recognition mode for diverse tRNA substrates by NSUN2

To test whether NSUN2 recognizes different tRNA substrates through a common mechanism, we determined the structures of NSUN2(C271A) in complex with tRNA^Lys^ and pre-tRNA^Leu^ at 3.93 Å and 3.30 Å resolution, respectively (Extended Data Figs. 5-6). Despite differences in length and maturation state, both substrates bind NSUN2 in a manner nearly identical to tRNA^Tyr^ (Figs. 3a, 4a, b; Extended Data Figs. 3a, 7a,b). Consistent with this shared binding topology, tRNA^Lys^ and pre-tRNA^Leu^ undergo conformational changes similar to those observed for tRNA^Tyr^ (Fig. 2a, b, d; Extended Data Fig. 7c-e).

Analysis of the higher-resolution pre-tRNA^Leu^ complex showed that most protein-RNA contacts are conserved relative to the tRNA^Tyr^ complex (Fig. 4c; Extended Data Fig. 8a). Differences are limited to a small number of base-specific interactions, such as recognition of A72 by N592 in the acceptor arm and recognition of A60 by R133 in the T-loop. In addition, N280, which recognizes U47 in tRNA^Tyr^, does not form a corresponding interaction with G47 in pre-tRNA^Leu^. These observations indicate that NSUN2 recognizes both mature and precursor tRNAs using a relatively conserved binding mode with limited sequence-specific adjustments. Unexpectedly, although pre-tRNA^Leu^ was previously reported to be methylated at C34 in the anticodon loop^13^, the pre-tRNA^Leu^ complex structure clearly shows the variable-loop cytosine C48 positioned in the catalytic pocket, with no density corresponding to C34 or the anticodon loop near the active site (Fig. 4b). To define the modification sites, we performed mutational analysis of C34 and C48. Single mutations at either position reduced methyltransferase activity, whereas the C34G/C48G double mutation abolished activity. Importantly, mutations at C48 caused a more pronounced loss of activity than corresponding mutations at C34, indicating that although NSUN2 can modify both sites, C48 represents the primary modification target in pre-tRNA^Leu^ (Fig. 4d).

The conservation of binding topology and target positioning prompted us to ask whether NSUN2 substrate selectivity reflects shared sequence features. We therefore compared the sequences of cytosolic tRNAs encoding all 20 amino acids together with pre-tRNA^Leu^. Only four tRNAs (tRNA^Arg^, tRNA^Asn^, tRNA^Cys^ and tRNA^Trp^) have not previously been reported as NSUN2 substrates. Sequence alignment revealed overall sequence conservation among tRNAs but no obvious consensus motif associated with NSUN2 modification (Extended Data Fig. 8b). Notably, among these four tRNAs, position 48 is uridine in three cases and cytidine only in tRNA^Cys^, raising the possibility that the presence of a cytidine at this position may influence NSUN2 recognition. Consistent with this idea, *in vitro* assays showed preferential methylation of tRNAs containing a cytidine at position 48 or 49 (Fig. 4e). Introduction of C48 into non-substrate tRNA^Asn^ and tRNA^Trp^ conferred NSUN2 activity, whereas mutation of C49 in tRNA^Arg^ reduced methylation efficiency. Finally, given the absence of stable interactions with the D-arm and anticodon arm, we asked whether NSUN2 requires only a minimal tRNA architecture for catalysis. Truncated “small tRNA” constructs containing the acceptor arm, T-arm and variable loop connected by flexible linkers were efficiently methylated by NSUN2, with the optimal construct retaining ∼88% of wild-type activity (Fig. 4f, g).

Together, these results demonstrate that NSUN2 recognizes diverse tRNA substrates through a conserved, structure-driven mechanism that relies on the acceptor arm, T-arm and variable loop, rather than on a fixed L-shaped tRNA architecture or strict sequence motifs. Although no global sequence consensus is evident, NSUN2 shows a strong preference for methylating cytidines at position 48 or 49, indicating that local nucleotide identity contributes to site selection within a largely structure-governed recognition mode.

### Structure-based screening of small-molecule inhibitors targeting NSUN2

To protect ongoing translational efforts, structure-based small-molecule screening and inhibitor characterization are not included in this preprint and will be disclosed separately.

## Discussion

NSUN2 is a central member of the NSUN family and is distinguished by its unusually broad substrate spectrum, including mature and precursor tRNAs, mRNAs and other non-coding RNAs^1^. Through these activities, NSUN2 participates in diverse biological processes such as cell proliferation and stress responses, and its dysregulation has been strongly linked to tumorigenesis^21^. However, the molecular basis by which a single RNA methyltransferase achieves such broad substrate recognition while maintaining site specificity has remained unclear. By determining structures of NSUN2 in the substrate-free state and in complex with multiple tRNA substrates, including tRNA^Tyr^, tRNA^Lys^ and pre-tRNA^Leu^, we establish a comprehensive structural framework for NSUN2-mediated tRNA m^5^C modification (Extended Data Fig. 9). Our data reveal a conserved recognition strategy in which NSUN2 primarily senses shared structural features of tRNA substrates rather than strict sequence motifs. A defining feature of this mechanism is the active induction of large-scale tRNA conformational rearrangements, which expose otherwise buried cytosines and precisely position them within the catalytic pocket. Such substrate remodeling represents a key regulatory step and provides a potential means for cells to modulate tRNA modification in response to physiological cues^28–31^.

Comparison with other NSUN family members highlights a fundamental divergence in substrate recognition strategies. NSUN6, whose structure and substrate recognition mechanism have been previously characterized, exhibits a highly restrictive mode of substrate selection that depends on precise recognition of local sequence and structural elements, including the CCA terminus, the target cytosine C72, and specific base pairs within the D-stem^9^. By contrast, although the catalytic MTase cores of NSUN2 and NSUN6 are highly conserved, NSUN2 contains an additional C-terminal PUA-like domain that enables a distinct, architecture-centered recognition logic (Extended Data Fig. 10a, b). Consistent with this difference, our structures show that NSUN2 does not primarily read sequence information; instead, it engages the conserved acceptor arm and T-arm scaffold of tRNA, destabilizes key long-range interactions at the tRNA elbow and inner corner, and actively remodels the substrate to expose the catalytic site. This structure-first, sequence-tolerant strategy explains how NSUN2 efficiently processes a large and diverse tRNA repertoire using a largely conserved interaction interface.

Induced tRNA remodeling has been observed in other modification systems^32^, such as the m^1^A58 methyltransferase Trm6-Trm61 complex, which flips a buried base into the active site through T-arm reorganization (Extended Data Fig. 10c, d). However, the mechanism uncovered here for NSUN2 is both structurally and functionally distinct. Unlike Trm6-Trm61, which requires an auxiliary tRNA-binding subunit, NSUN2 integrates substrate capture and catalysis within a single polypeptide. Moreover, NSUN2 induces more extensive conformational changes, affecting not only the T-arm but also the acceptor arm and variable loop, ultimately allowing target cytosine to access the catalytic center. This pronounced remodeling accounts for the high conformational flexibility of the D-arm and anticodon arm observed in our structures and underscores NSUN2’s exceptional substrate adaptability.

The structural insights gained here also have direct implications for therapeutic targeting of NSUN2. NSUN2 is frequently overexpressed or functionally altered in cancers, where enhanced tRNA m^5^C modification is thought to support elevated translational demand. Unlike SAM-competitive inhibitors that target the conserved cofactor-binding site and frequently display poor specificity across methyltransferases^33,34^, our structures identify a druggable pocket at the MTase-PUA-like domain interface that is distal to the catalytic center and functionally coupled to substrate RNA binding. Using structure-based virtual screening, we identified the small molecule X as a reversible inhibitor of NSUN2. Mechanistically, X disrupts NSUN2-RNA interactions rather than cofactor binding, selectively reducing tRNA m^5^C levels in cells without altering global RNA methylation or NSUN2 protein abundance. Given the established role of tRNA m^5^C in maintaining translational efficiency in rapidly proliferating cells, inhibition of NSUN2 provides a plausible molecular basis for the observed suppression of cancer cell growth. Although X represents an early-stage lead, its identification establishes proof of principle that NSUN2 can be selectively targeted through non-catalytic interfaces.

In summary, this study defines a conserved yet highly dynamic mechanism by which NSUN2 recognizes and remodels tRNA substrates to achieve broad specificity and precise catalysis. By coupling structural analysis with functional and biochemical approaches, we clarify the molecular basis of NSUN2-mediated tRNA methylation and provide a structural foundation for developing selective NSUN2 inhibitors, opening new avenues for targeting RNA modification pathways in diseases.

## Methods and Materials

### Cloning, Expression, and Purification of NSUN2 Constructs

The human NSUN2 sequence (UniProtKB: Q08J23) was codon-optimized for expression in *Escherichia coli* (*E. coli*). The gene was cloned into a pET-28 vector, incorporating an N-terminal 6×His tag. The recombinant plasmid was transformed into *E. coli* BL21 (DE3) cells, which were initially cultured overnight at 37°C in LB medium supplemented with kanamycin. The culture was subsequently scaled up until the optical density at 600 nm (OD600) reached 0.8. Protein expression was induced with 0.5 mM isopropyl β-D-1-thiogalactopyranoside (IPTG) followed by incubation at 16°C for 20-24 h. Cells were harvested by centrifugation at 5000 × g for 10 min at 4°C. The cell pellet was resuspended in Binding Buffer (20 mM Tris, 1 M NaCl, pH 7.5) and lysed using a high-pressure homogenizer. The lysate was cleared by centrifugation at 10,000 × g for 30 min at 4°C, and the resulting supernatant was subjected to affinity purification using a Ni-NTA column (GE Healthcare). Further purification was performed via size-exclusion chromatography (SEC) using a HiLoad 16/60 Superdex 200 column (GE Healthcare). The purified protein was stored in a final buffer of 20 mM Tris and 200 mM NaCl (pH 7.5). NSUN2 mutants were generated via site-directed mutagenesis. Linearized fragments containing the desired mutations were amplified by PCR, treated with a MutanBEST Kit (Takara) for phosphorylation and blunting, and circularized using Solution I. The expression and purification protocols for the mutant proteins were identical to those used for the wild-type protein.

### Cloning, *In Vitro* Transcription, and Purification of RNAs

To facilitate *in vitro* transcription, a T7 promoter and leader "G" nucleotides were appended upstream of the target DNA sequence, which was then cloned into a pUC19 vector. Double-stranded DNA templates were amplified via PCR. To minimize N+1 transcriptional heterogeneity common with T7 RNA polymerase, the reverse primers were designed with 2′-O-methyl modifications on the two penultimate 5′-terminal nucleotides. Transcription was performed in a reaction mixture containing transcription buffer [40 mM Tris-HCl, 1 mM spermidine, 0.01% (v/v) Triton X-100, pH 8.1], 10 mM DTT, 5 mM NTPs, 30 mM MgCl2, 0.3 mg/mL T7 RNA polymerase, and 10 ng/µL DNA template. The reaction was incubated at 37°C for 4 h and terminated by heat inactivation at 72°C for 20 min. EDTA was added to a final concentration of 50 mM to chelate magnesium ions. RNA was precipitated with three volumes of ethanol at -80°C overnight. The pellet was resuspended in DEPC-treated water and mixed with an equal volume of 2× denaturing loading buffer. The mixture was loaded onto a 10% urea-denaturing polyacrylamide gel to purify RNA. The target band was visualized by ultraviolet shadowing, excised and incubated overnight in a gel elution buffer to elute the RNA. The recovered RNA was ethanol-precipitated, resuspended in DEPC-treated water and stored at -80°C. RNA mutants were constructed and purified following the same procedures as the wild-type RNA.

### *In Vitro* Methyltransferase Activity Assay

Methyltransferase activity was assayed in a 40 µL reaction volume. The mixture consisted of 50 mM Tris-HCl (pH 8.0), 50 mM ammonium acetate (NH4Ac), 3 mM MgCl2, 1 mM DTT, 0.5 µM recombinant protein, 1 µM RNA substrate, and 1 µL of 0.5 µCi/µL ³H-labeled S-adenosylmethionine (³H-SAM; PerkinElmer). The reaction was quenched by adding 100 µL of water-saturated phenol. Following vigorous vortexing and centrifugation (13,000 × g, 10 min, 4°C), the upper aqueous phase was recovered. RNA was further purified using a chloroform:isoamyl alcohol (24:1) extraction and precipitated with pre-chilled absolute ethanol at -40°C for 1 h. The RNA pellet was collected, washed with 75% ethanol, and air-dried. The dried pellet was resuspended in 10 µL of DEPC-treated water and transferred to a scintillation vial containing 5 mL of scintillation fluid (Ultima Gold™, PerkinElmer). Radioactivity was measured using a liquid scintillation counter (Beckman Coulter LS6500) to quantify ³H-methyl incorporation. In the small-molecule enzyme inhibition assay, compounds were added to the reaction system at a final concentration of 50 μM, with an equivalent volume of DMSO used as the control. To determine IC50 values, equal volumes of compounds at different concentrations were added to the reactions, and IC50 values were calculated using GraphPad Prism software.

### Electrophoretic Mobility Shift Assay (EMSA)

In the EMSA, five samples were set up, each with a reaction volume of 10 µL. Each reaction contained RNA at a final concentration of 0.5 µM. The first reaction served as a protein-free control, while increasing concentrations of the protein were added to the remaining four reactions. All mixtures were adjusted to a final volume of 10 µL using an incubation buffer consisting of 20 mM Tris (pH 7.8), 150 mM NaCl, and 5 mM MgCl2, followed by incubation at 4 °C for 1 h. After incubation, 2.5 µL of glycerol loading buffer was added to each sample. The samples were then resolved on a 6 % polyacrylamide gel at a constant voltage of 120 V for approximately 20 min. Subsequently, the gel was stained with GelRed for 1 min and visualized using a gel documentation system. In the small-molecule competitive EMSA, compounds at a final concentration of 100 μM or an equal volume of DMSO were added to protein-RNA complex samples, in which the final concentrations of protein and RNA were 1 μM and 100 nM, respectively. Detection was performed as described above.

### Fluorescence Polarization (FP) Binding Assay

5’-Carboxyfluorescein (FAM)-labeled tRNA (Icor Bioscience) was dissolved in DEPC-treated water to a 100 µM stock concentration. Prior to the assay, the RNA was refolded by heating at 95°C for 3 min followed by slow cooling to room temperature. The optimal RNA concentration for the FP assay was determined to be 40 nM. Serial dilutions of the protein (up to 40 µM) were prepared and mixed with 40 nM FAM-tRNA in 96-well black microplates (Greiner). After a 10 min incubation on ice, FP was measured using a CLARIOstar microplate reader. Dissociation constants (*K*D) were determined by fitting the data to a 1:1 binding stoichiometry using GraphPad Prism. The following equation was utilized^35^:

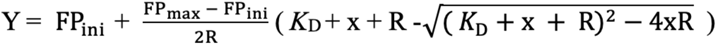

where Y is the total FP value, FPmin is the initial polarization value (signal of free RNA), FPmax is the maximum polarization value (signal of the fully bound complex), *K*D is the dissociation equilibrium constant, R is the total RNA concentration, and x is the total protein concentration. In the small-molecule inhibition assay of NSUN2-tRNA binding, compounds were added to each well at a final concentration of 50 μM. Detection and data analysis were performed as described above.

### Crystallization and Structural Determination

Purified NSUN2 (10 mg/mL) was mixed with S-adenosyl-L-homocysteine (SAH) at a 1:3 molar ratio. Crystals were grown via the sitting-drop vapor diffusion method in a reservoir solution containing 25% (w/v) PEG 3350, 0.2 M Lithium Sulfate, and 0.1 M HEPES (pH 7.5). X-ray diffraction data were collected at the Shanghai Synchrotron Radiation Facility (SSRF) beamline BL19U1 and processed using the XDS package^36^. Data scaling and space group analysis were performed using Aimless from the CCP4 suite^36,37^. Initial phases were determined by molecular replacement using MolRep, employing the crystal structure of human NSUN6 (PDB ID: 5WWQ) as a search model^38,39^. The structural model was refined using Refmac5^40^ and Phenix^41^, with manual model building and adjustment performed in Coot^42^.

### Complex Reconstitution for Cryo-EM Analysis

To prepare the complex for cryo-EM analysis, the RNA was first refolded *in vitro* by heating at 95°C for 5 min, followed by immediate cooling on ice. The NSUN2-RNA complex was then assembled by incubating the purified NSUN2 protein, the refolded tRNA, and S-adenosyl-methionine (SAM) for 1 h on ice, maintaining a molar ratio of 1:1.5:3. The resulting incubation mixture was purified via size-exclusion chromatography (SEC) using a HiLoad 10/300 Superdex 200 Increase column (GE Healthcare), which was pre-equilibrated with a buffer containing 20 mM Hepes (pH 7.5), 200 mM NaCl, and 2 mM MgCl2. Peak fractions corresponding to the stable complex were collected and concentrated to approximately 1 mg/mL for subsequent cryo-EM grid preparation.

### Cryo-EM Grid Preparation and Data Collection

Cryo-EM grids (Quantifoil R2/1 or Quantifoil R1.2/1.3, 200-mesh) were glow-discharged using a plasma cleaning system with a 50-second exposure and a 15-second hold time. Grids were then transferred into a Vitrobot maintained at 4°C and 100% humidity. A 4 μL aliquot of the sample was applied to each grid, followed by a 3.5 s blot time and 5 s wait time, before the grids were rapidly plunge-frozen in liquid ethane. Initial grid screening was performed on a Glacios cryo-electron microscope (Thermo Fisher Scientific) operating at 200 kV. High-resolution data collection was subsequently carried out using a Titan Krios cryo-electron microscope (Thermo Fisher Scientific) at 300 kV, with a nominal magnification of 130,000 × (yielding a calibrated pixel size of 0.93 Å or 0.97 Å). Micrographs were acquired using EPU software equipped with a Falcon 4i direct electron detector. A total of 4024 movie stacks for NSUN2-tRNA^Tyr^, 5548 movie stacks for NSUN2-tRNA^Lys^, and 4250 movie stacks for NSUN2-pre-tRNA^Leu^ were collected within a defocus range of -1.5 to -2.5 μm.

### Cryo-EM Data Processing

Raw movie stacks were processed using the cryoSPARC software package^43^. Motion correction was performed via the Patch Motion Correction algorithm, and the Contrast Transfer Function (CTF) parameters for each micrograph were estimated using Patch CTF Estimation. Particle picking was executed using the Template Picker in cryoSPARC. Following 2D classification, a subset of 100,000 high-quality particles was selected for ab initio reconstruction, followed by 3D heterogeneous refinement. Particles from the most promising 3D classes were initially refined using non-uniform refinement, followed by reference-based motion correction and subsequent rounds of non-uniform and local refinement to further improve the resolution. The final map resolutions were determined based on the gold-standard Fourier Shell Correlation (FSC) 0.143 criterion. Local resolution maps were generated within cryoSPARC using the "Local Resolution Estimation" tool. Detailed processing workflows and statistics are provided in Extended Data Figs. 2, 5, 6 and Extended Data Tables 2-4.

### Model Building and Refinement

For the NSUN2-pre-tRNA^Leu^-SAM complex, the crystal structure of NSUN2-SAH was utilized as the initial search model, while the pre-tRNA^Leu^ density was modeled *de novo*. The atomic model was iteratively optimized through manual adjustment in Coot and automated refinement using phenix.real_space_refine. The refined coordinates of the NSUN2-pre-tRNA^Leu^-SAM complex subsequently served as the starting template for building the NSUN2-tRNA^Tyr^-SAM structure, which was optimized using the same computational pipeline. Finally, the refined NSUN2-tRNA^Tyr^-SAM structure was used as the initial model for the NSUN2-tRNA^Lys^-SAM complex, followed by a final round of optimization in Coot and phenix to ensure optimal stereochemistry and agreement with the cryo-EM density. The quality of final models was evaluated using MolProbity^44^. Model statistics are provided in Extended Data Tables 2-4. Figures were made with Chimera^45^ or ChimeraX^46^. Secondary structure diagrams were prepared using RiboDraw, followed by manual optimization (https://github.com/ribokit/RiboDraw).

### Virtual Screening

The ProteinsPlus web server (https://proteins.plus/) was used to analyze pocket properties of the NSUN2-SAH structure, including pocket volume, depth, number of hydrogen bond donors and acceptors, and hydrophobicity. The screening pocket was selected based on the overall scoring. A laboratory in-house library of 785 low-molecular-weight conventional drugs, with molecular weights of approximately 300 Da, was used for virtual screening. Three-dimensional structures of the small molecules were generated using RDKit^47^, followed by 500 steps of conformational optimization with the MMFF94S^48^ force field. Binding free energies between the compounds and the selected receptor pocket were calculated using Smina and ranked accordingly. Compounds with the lowest predicted binding free energies were selected for further *in vitro* enzymatic activity assays.

### Cell Culture

Human gastric cancer cell line HGC27 was cultured in Dulbecco’s Modified Eagle Medium (DMEM; Gibco) supplemented with 10% fetal bovine serum and 1% penicillin-streptomycin. Cells were maintained at 37°C in a humidified incubator with 5% CO2.

### Cell Proliferation Assay

Gastric cancer cell line HGC27 was seeded into 96-well plates at a density of approximately 5,000 cells per well and cultured at 37°C in a humidified incubator with 5% CO2. On the following day, cells were treated with the indicated compounds or DMSO for 48 h. Cell proliferation was assessed using the Cell Counting Kit-8 (CCK-8; Abbkine, Inc. China) according to the manufacturer’s instructions. Optical density (OD) was measured at 450 nm using a CLARIOstar microplate reader. IC50 values were calculated using GraphPad Prism software.

### Western Blot

Cells were lysed on ice using RIPA lysis buffer (Beyotime), followed by the addition of SDS loading buffer. Total protein lysates were separated by SDS-PAGE and transferred onto nitrocellulose (NC) membranes. After washing three times with 0.1% TBST, the membranes were sequentially incubated with primary antibodies and the horseradish peroxidase-conjugated anti-mouse secondary antibodies (Thermo, 31430). Protein bands were visualized using a chemiluminescent HRP substrate (Millipore, USA) and detected by autoradiography using LAS4000mini (GE bio. Inc.). GAPDH (Proteintech, HRP-60004) was used as a loading control. The NSUN2 protein was detected using the monoclonal antibody (Proteintech, 66580-1-lg).

### RNA m^5^C Dot Blot Assay

Total RNA and small RNA (< 200 nt) were extracted from DMSO- or compound-treated cells using the MolPure Cell/Tissue Total RNA Kit (Yeasen) and the RNA Clean & Concentrator Kit (Zymo Research, R1013), respectively. RNA samples at different concentrations were spotted onto Hybond N^+^ nylon membranes (Beyotime). After UV crosslinking for 5 min, membranes were blocked with TBST containing 5% nonfat milk and then sequentially incubated with a mouse anti-m^5^C primary antibody (Proteintech, 68301-1-lg) and the corresponding anti-mouse secondary antibody. After three washes with TBST, dot blot signal intensity was detected by autoradiography.

## Acknowledgments

We thank the Cryo-EM Centers at the University of Science and Technology of China (USTC), Huazhong University of Science and Technology (HUST) and Anhui Medical University (AHMU) for their assistance with the experiments. We also thank Dr. Ke Ruan’s laboratory for providing the small-molecule repurposed drug library, Dr. Xiaofeng Zhang’s laboratory for providing multiple tRNA template plasmids, Dr. Zhenye Yang’s laboratory for providing the HGC27 cell line and Ming Li for assistance with small-molecule complex prediction. This work was supported by the National Natural Science Foundation of China (92581123 and 32371345 to K.Z., 32371310 to Q.G., 32301044 and 32471301 to S.L.), the National Key R&D Program of China (2022YFA1302700 to K.Z.), the Strategic Priority Research Program of the Chinese Academy of Sciences (XDB0490000 to K.Z.), the Center for Advanced Interdisciplinary Science and Biomedicine of IHM (QYPY20220019 to K.Z. and QYPY20220004 to Q.G.), the International Partnership Program of Chinese Academy of Sciences (123GJHZ2024080FN to S.L.), the China Postdoctoral Science Foundation (General Program, 2024M763148 to Q.H. and 2025M782790 to W.Y.), and the China Postdoctoral Science Foundation-Anhui Joint Support Program ( 2024T005AH to Q.H.).

## Author Contributions

K.Z., Q.G., and S.L. conceived the study and designed the experiments. Q.H., W.Y., Y.Y., and R.Y. prepared the samples, performed biochemical experiments, and carried out cryo-EM sample preparation, screening, data collection, image processing. Y.Z. and L.D. helped with sample preparation. L.W. performed the virtual screening. F.L. solved the crystal structure. K.Z. conducted final cryo-EM structure determination. W.Y. and S.L. built and refined the cryo-EM models. Q.H., W.Y., K.Z., Q.G., and S.L. analyzed the data. Q.H., W.Y., K.Z., Q.G., and S.L. wrote and edited the manuscript.

## Competing Interests

Authors declare that they have no competing interests.

## Data and Materials Availability

Cryo-EM structures and atomic models generated in this study have been deposited in the wwPDB OneDep System under EMD accession codes EMD-68138, EMD-68140, EMD-68111 and PDB ID codes under accession codes 22AV, 22AX, 22ZH, respectively. Coordinate and structure factor for NSUN2-SAH has been deposited in the wwPDB OneDep System under accession code 22BM.

## Notes

### Competing Interest Statement

The authors have declared no competing interest.

