## Supplementary Information for "Structural insights into the recognition and catalysis of tRNA by human NSUN2"

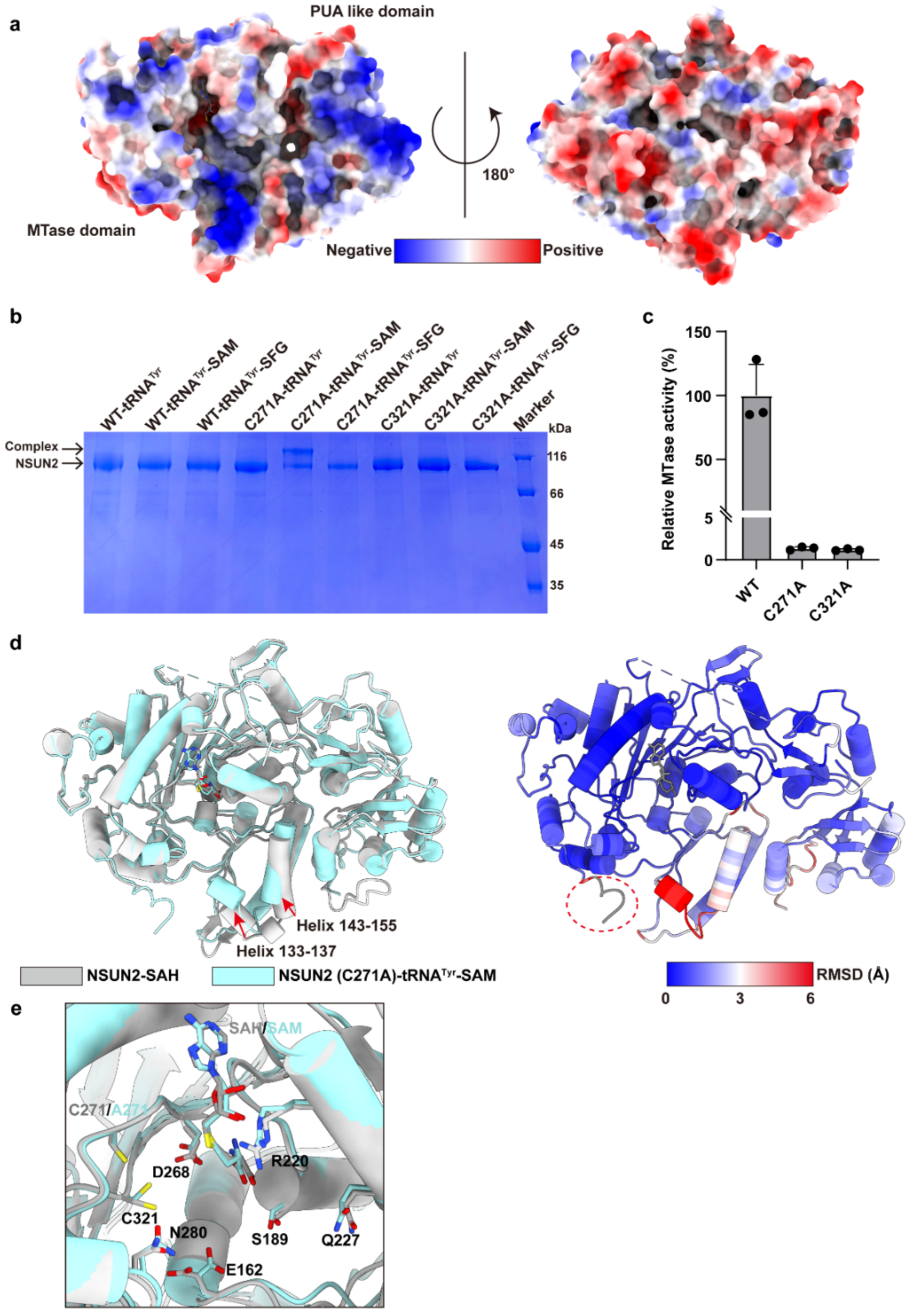


**Extended Data Fig. 1: Structural snapshots of NSUN2 before and after tRNA binding.**

**a.** Electrostatic surface representation of the NSUN2-SAH. **b.** SDS-PAGE analysis of complexes formed between tRNA and wild-type or mutant NSUN2. SFG (Sinefungin) is an analog of S-adenosyl-methionine (SAM) **c.** Comparison of relative activities of wildtype and variant NSUN2 methylating tRNA$^{Tyr}$. Data are shown as the mean ± SD (n = 3 independent experiments). **d.** Structural comparison of the NSUN2 in NSUN2-SAH and NSUN2-tRNA$^{Tyr}$-SAM complexes (left). The NSUN2 in the NSUN2-tRNA$^{Tyr}$-SAM complex is colored by C-alpha RMSD in ChimeraX (right). **e.** Active-site residue comparison between NSUN2-SAH and NSUN2-tRNA$^{Tyr}$-SAM complexes.

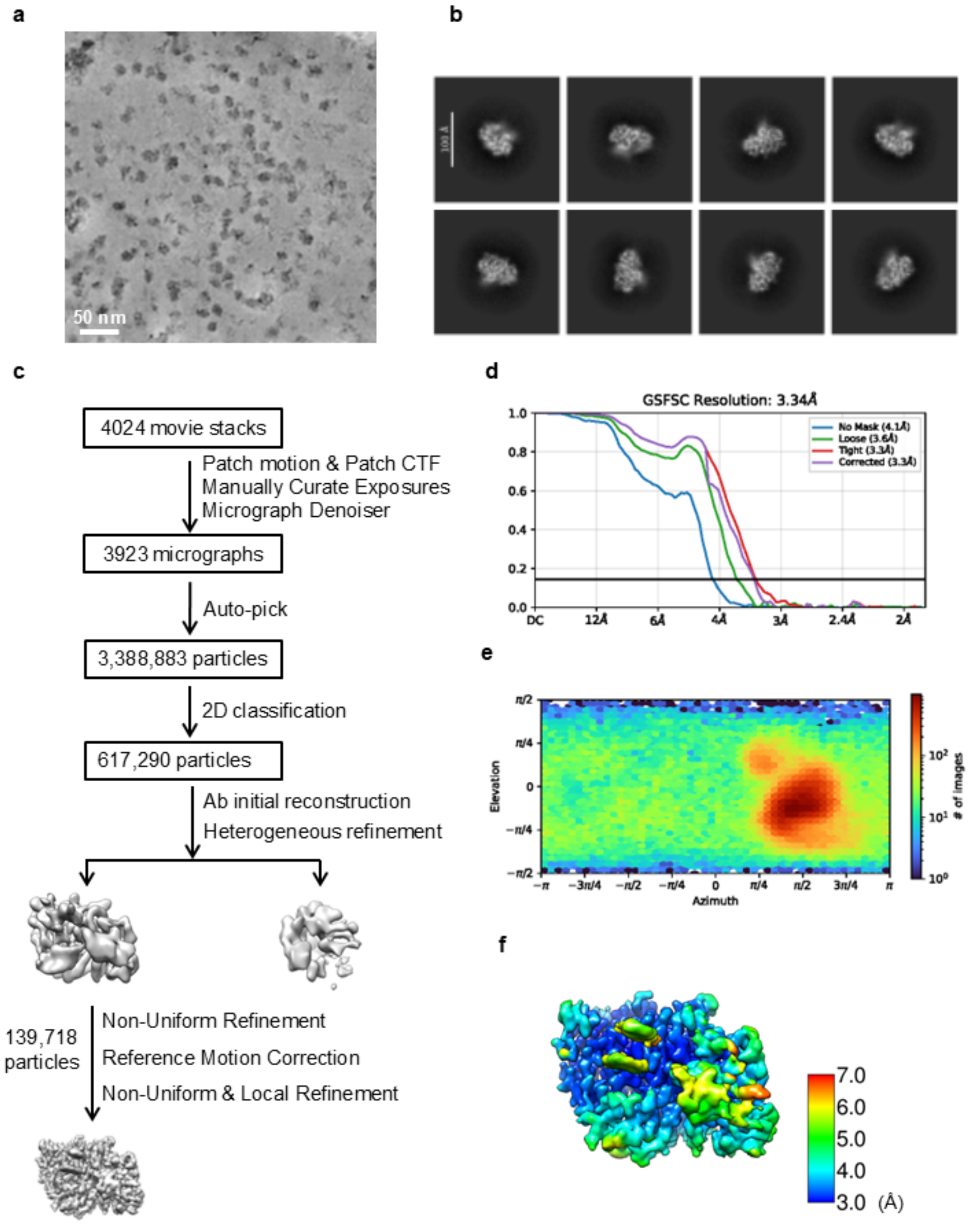


**Extended Data Fig. 2: Single-particle cryo-EM analysis of the NSUN2 (C271A) with tRNA$^{Tyr}$ substrate.**

**a.** Representative motion-corrected and denoised cryo-EM micrograph. **b.** Reference-free 2D class averages. **c.** Workflow of the data processing. **d.** Gold standard FSC plot for the 3D reconstruction, calculated in cryoSPARC. **e.** Euler angle distribution of the particle images. **f.** Resolution map for the final 3D reconstruction.

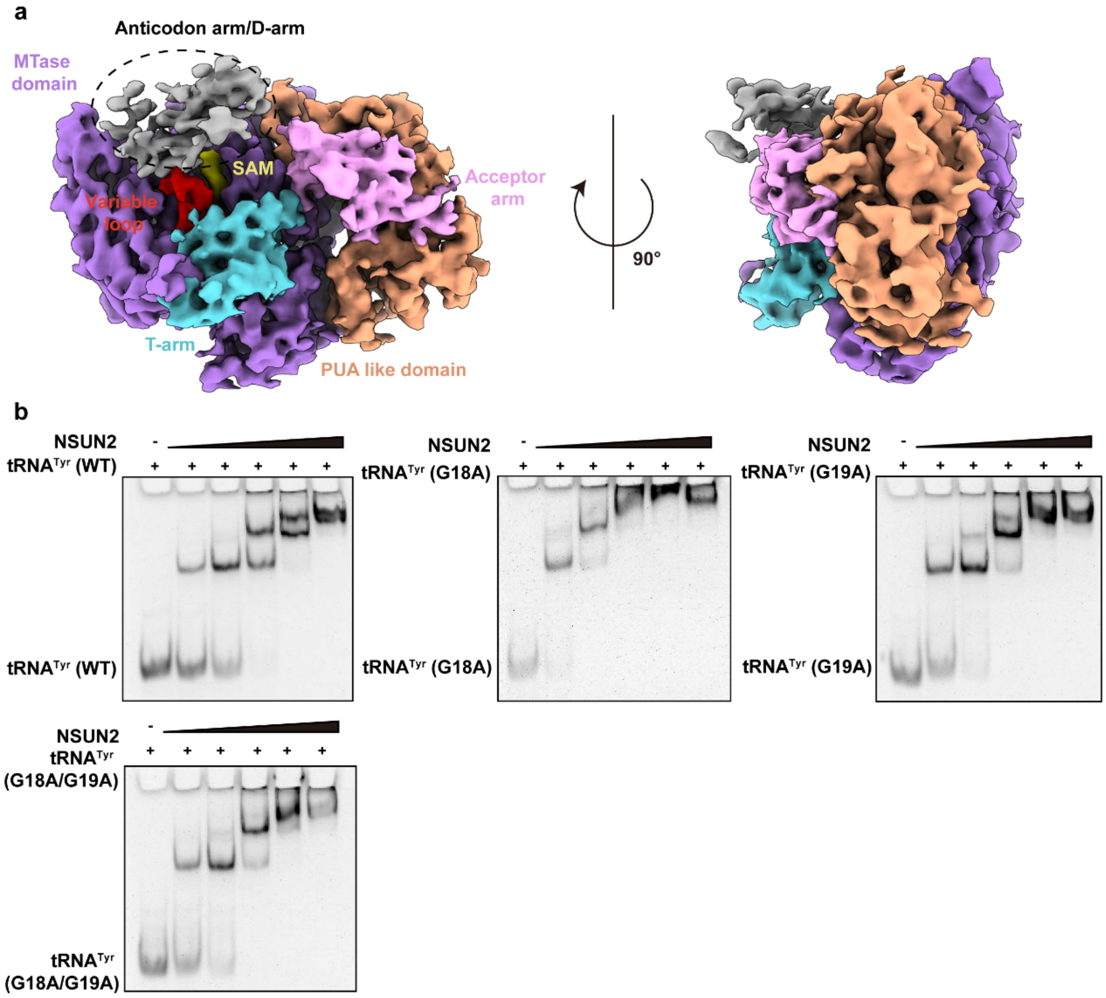


**Extended Data Fig. 3: L-shaped tRNA integrity is dispensable for NSUN2 binding.** **a.** The cryo-EM map of the NSUN2-tRNA$^{Tyr}$-SAM, showing the poorly resolved Anticodon arm/D-arm region. **b.** EMSA analysis of the binding affinity between NSUN2 and wild-type or mutant tRNAs.

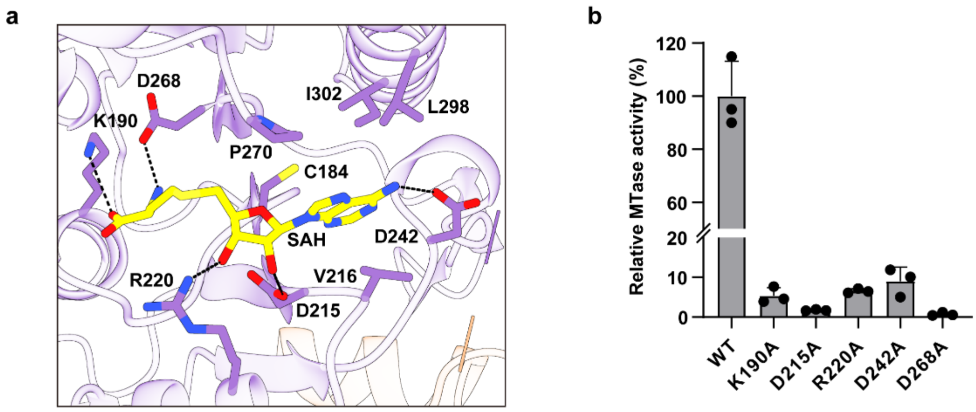


**Extended Data Fig. 4: Interactions between NSUN2 and SAH.**

**a.** Detailed view of NSUN2-SAH interactions. **b.** Comparison of relative activities of wildtype and variant NSUN2 methylating $tRNA^{Tyr}$. Data are shown as the mean ± SD (n = 3 independent experiments).

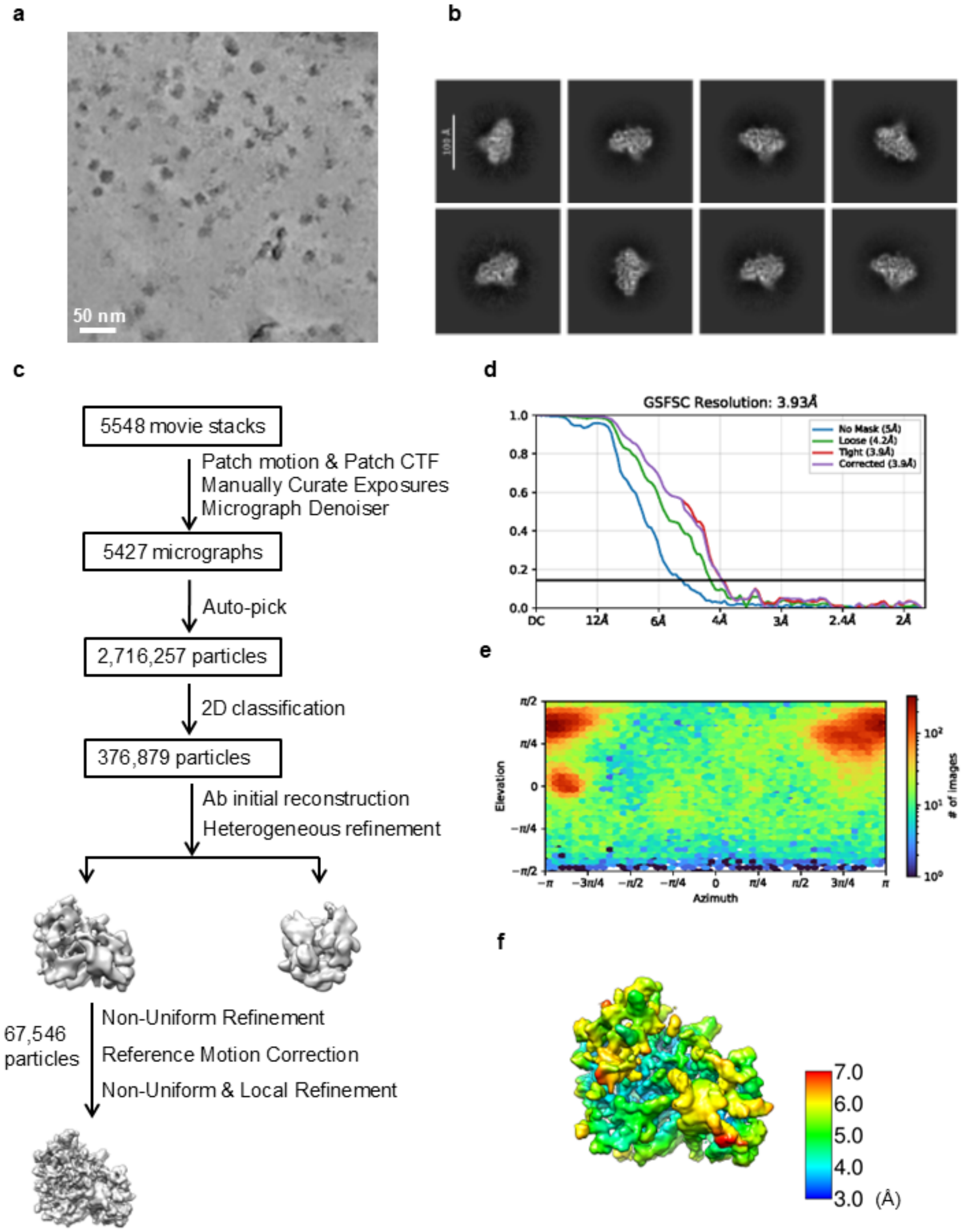


**Extended Data Fig. 5: Single-particle cryo-EM analysis of the NSUN2 (C271A) with tRNA$^{Lys}$ substrate.**

**a.** Representative motion-corrected and denoised cryo-EM micrograph. **b.** Reference-free 2D class averages. **c.** Workflow of the data processing. **d.** Gold standard FSC plot for the 3D reconstruction, calculated in cryoSPARC. **e.** Euler angle distribution of the particle images. **f.** Resolution map for the final 3D reconstruction.

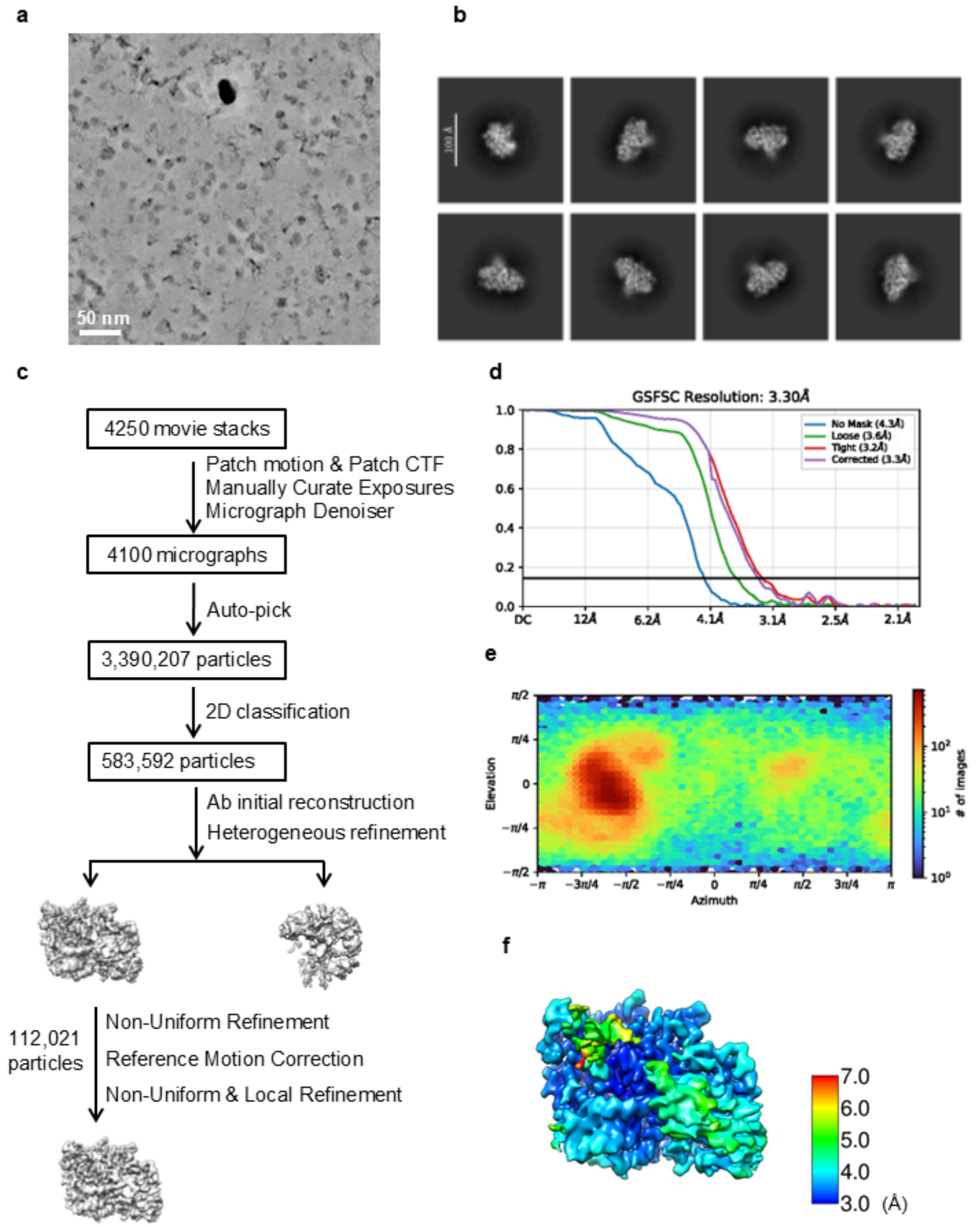


**Extended Data Fig. 6: Single-particle cryo-EM analysis of the NSUN2 (C271A) with pre-tRNA$^{Leu}$ substrate.**

**a.** Representative motion-corrected and denoised cryo-EM micrograph. **b.** Reference-free 2D class averages. **c**. Workflow of the data processing. **d.** Gold standard FSC plot for the 3D reconstruction, calculated in cryoSPARC. **e.** Euler angle distribution of the particle images. **f.** Resolution map for the final 3D reconstruction.

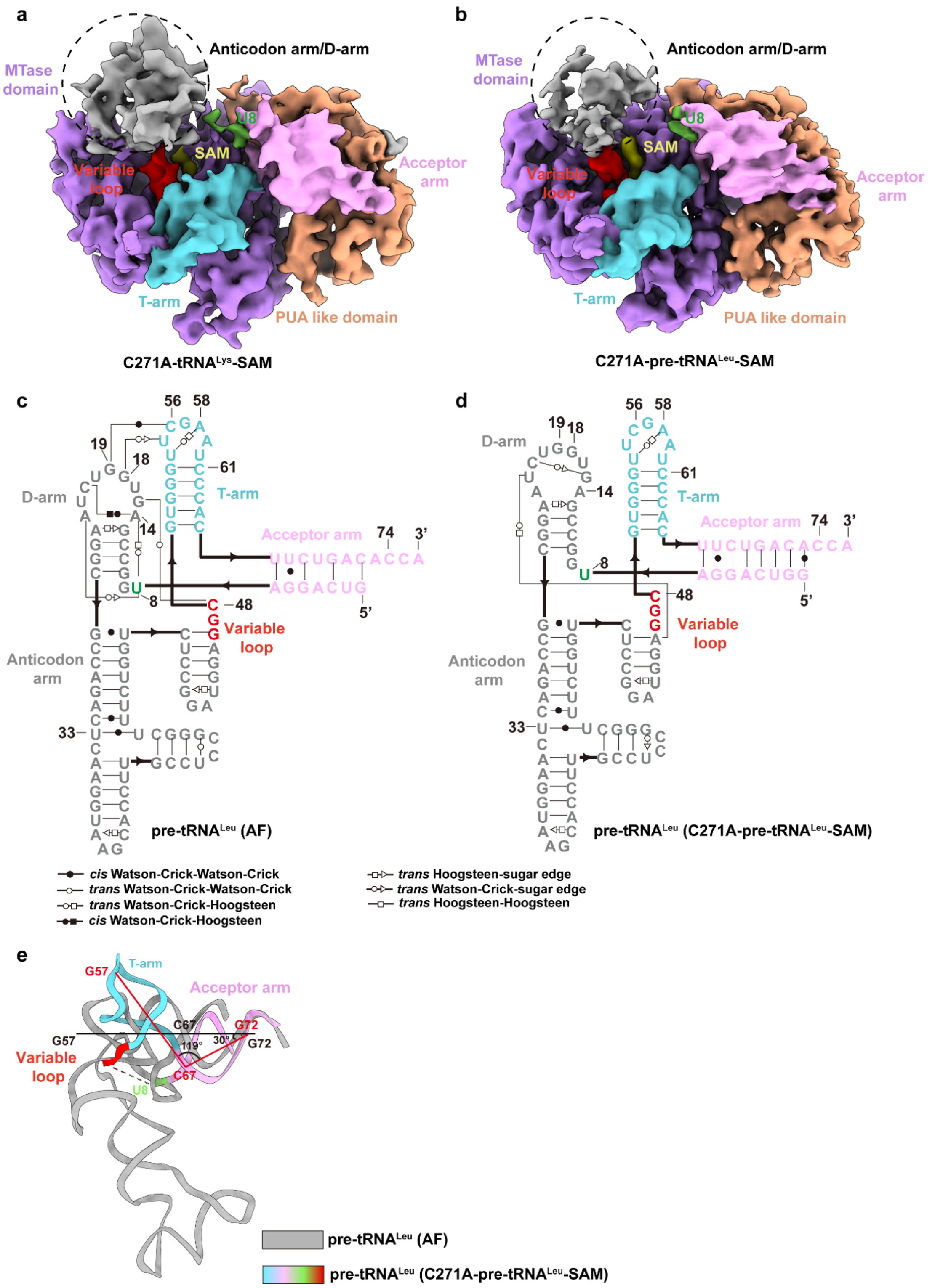


**Extended Data Fig. 7: Structural analysis of the NSUN2-tRNA$^{Lys}$-SAM and NSUN2-pre-tRNA$^{Leu}$-SAM complex.**

**a-b.** The cryo-EM map of the NSUN2-tRNA$^{Lys}$-SAM (a) and NSUN2-pre-tRNA$^{Leu}$-SAM (b). **c-d.** Secondary structure schematic of pre-tRNA$^{Leu}$ predicted by AlphaFold (c) and that observed in the NSUN2-pre-tRNA$^{Leu}$-SAM complex (d). **e.** Comparison between the pre-tRNA$^{Leu}$ predicted by AlphaFold and observed in the NSUN2-pre-tRNA$^{Leu}$ -SAM complex.

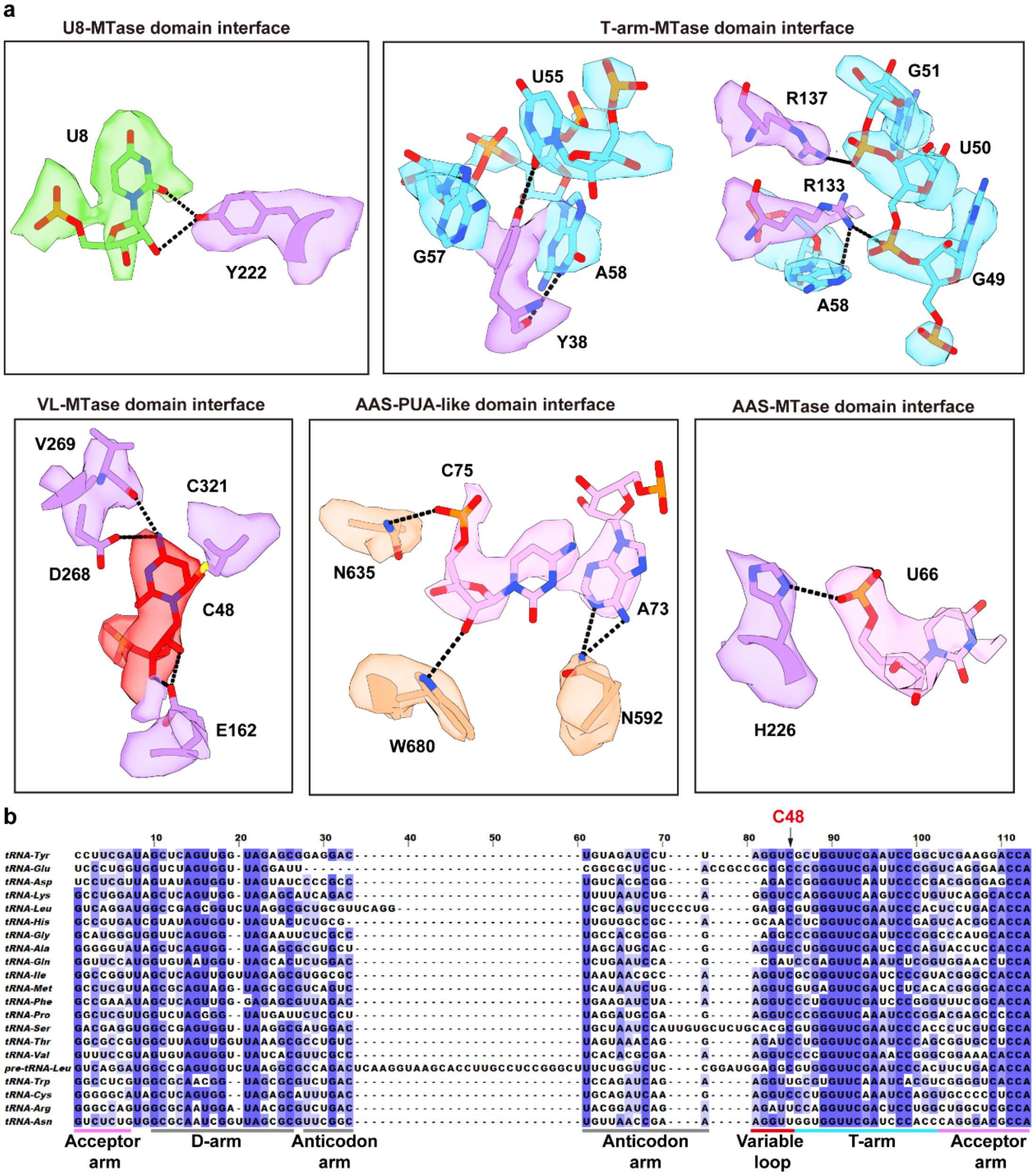


**Extended Data Fig. 8: Structural analysis of NSUN2-pre-tRNA$^{Leu}$ interactions**

**a.** Detailed view of NSUN2-pre-tRNA$^{Leu}$ interactions. **b.** Sequence alignment of tRNAs for all 20 amino acids and pre-tRNA$^{Leu}$.

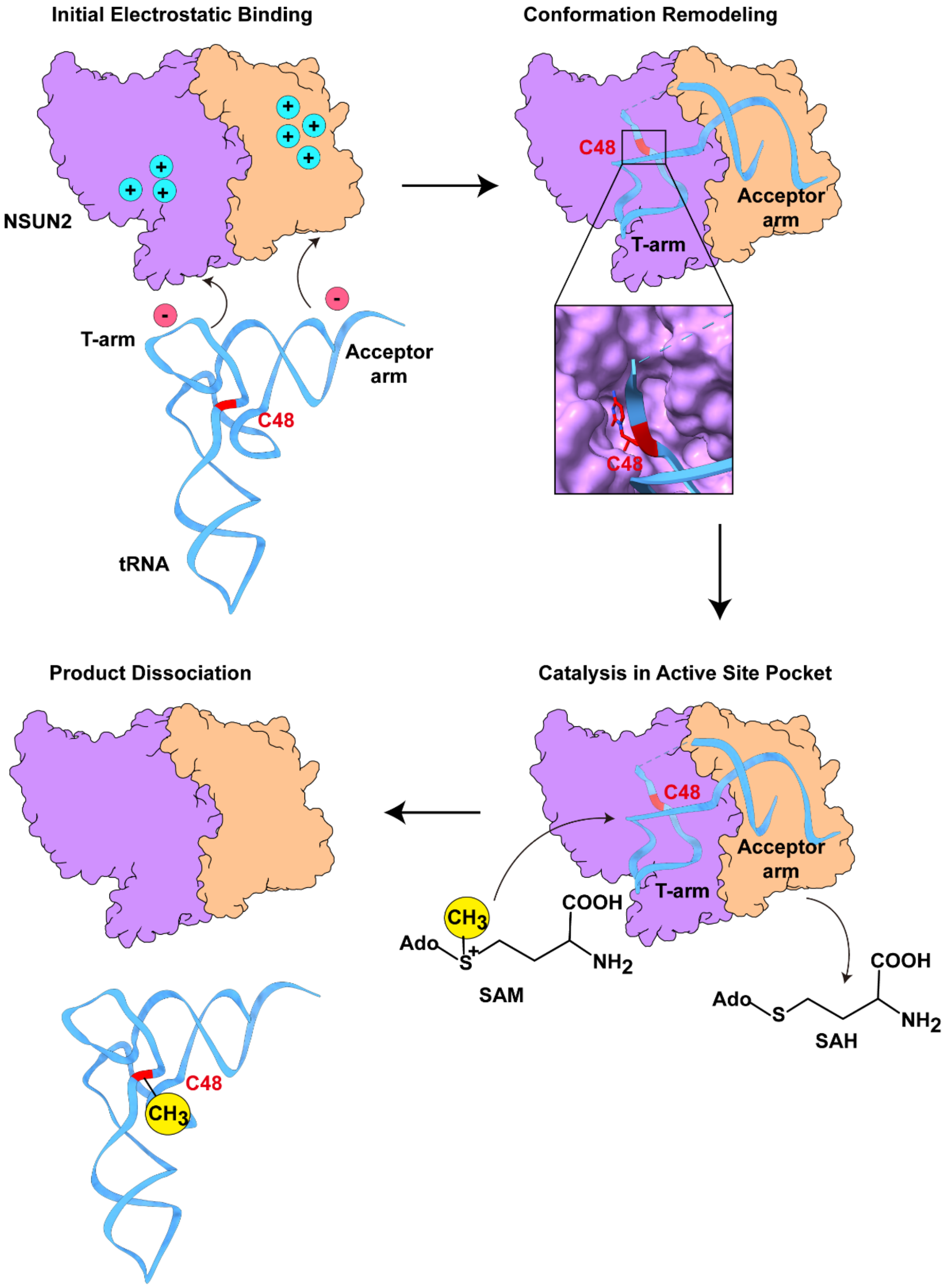


**Extended Data Fig. 9: Proposed mechanistic model of NSUN2-mediated tRNA methylation.**

Step 1. Initial electrostatic anchoring. NSUN2 initially captures the tRNA through a bipartite, structure-sensing basic surface via electrostatic interactions. Step 2. Conformation Remodeling. NSUN2 induces extensive tRNA conformational rearrangements by applying mechanical force to the tRNA elbow and inner region. Step 3. Catalysis in Active Site Pocket. Once C48 is precisely positioned in the active site, SAM donates a methyl group to generate m$^5$C48 and SAH. Step 4. Product dissociation. Following catalysis, the methylated tRNA is released, accompanied by restoration of its native conformation, completing the catalytic cycle.

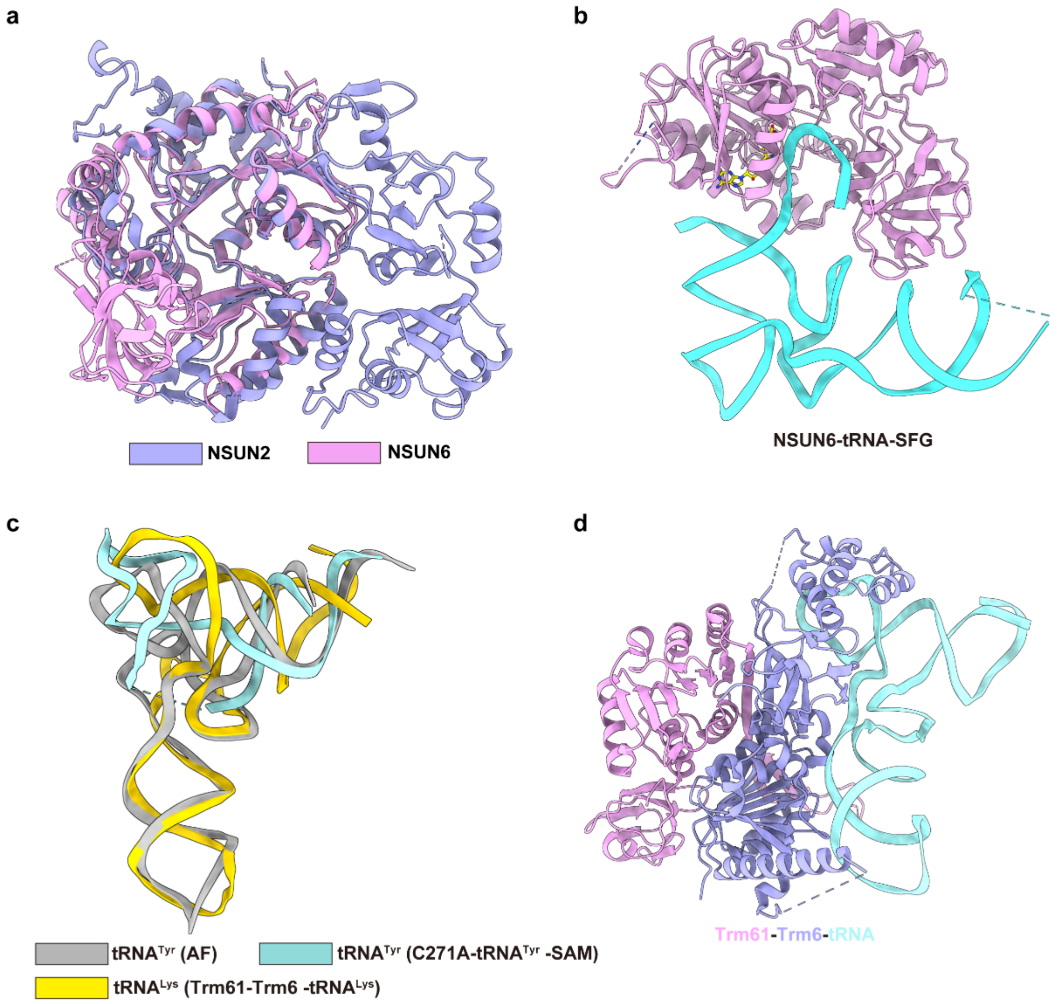


**Extended Data Fig. 10: Structural comparison of NSUN2 with other methyltransferases.**

**a.** Structural comparison of NSUN2 with NSUN6. **b.** Structure of NSUN6-tRNA-SFG. **c.** Structural alignment of tRNA in the predicted tRNA, NSUN2-$tRNA^{Tyr}$ complex, and the Trm61-Trm6 complex. **d.** Structure of Trm61-Trm6-tRNA.

**Extended Data Table 1. X-Ray data collection, processing, and model validation of the NSUN2-SAH.**

| | NSUN2-SAH |
|---|---|
| **Data collection** | |
| Beamline | BL19U1, SSRF |
| Space group | P1 |
| Wavelength (Å) | 0.979 |
| Resolution (Å) | 19.90-2.50 (2.59-2.50)[a] |
| Cell parameters | |
| a, b, c (Å) | 77.98, 88.35, 90.76 |
| α, β, γ (°) | 104.72, 99.57, 112.50 |
| Unique reflections | 69683 (6679) |
| Completeness (%) | 96.10 (92.98) |
| Redundancy | 3.6 (3.4) |
| I/σI | 8.51 (1.14) |
| $R_{merge}$ (%) | 14.95 (103.10) |
| $CC_{1/2}$ | 0.987 (0.583) |
| $CC^*$ | 0.997 (0.858) |
| **Refinement** | |
| $R_{work}$ (%) | 21.14 |
| $R_{free}$ (%) | 25.64 |
| No. of atoms | |
| Protein | 13848 |
| SAH | 26 |
| Water | 120 |
| Average B factors (Å$^2$) | |
| Protein | 53.88 |
| SAH | 42.60 |
| Water | 43.81 |
| Root mean square deviations | |
| Bond lengths (Å) | 0.002 |
| Bond angles (°) | 0.51 |
| Ramachandran plot | |
| Favored (%) | 98.23 |
| Allowed (%) | 1.77 |
| Disallowed (%) | 0 |

**Table note:** [a]Values in parentheses indicate those corresponding to the highest-resolution shell.

**Extended Data Table 2. Cryo-EM data collection, processing, and model validation of the NSUN2- tRNA$^{Tyr}$-SAM complex.**

| | NSUN2- tRNA$^{Tyr}$-SAM complex |
|---|---|
| **Data collection and processing** | |
| Microscope | Titan Krios G4 |
| Voltage (kV) | 300 |
| Camera | Falcon4i |
| Grid type | R 1.2/1.3 Quantifoil copper grid (200 mesh) |
| Magnification | 130,000× |
| Pixel size (Å) | 0.93 |
| Total exposure ($e^-$/Å$^2$) | 50 |
| Exposure time (s) | 5.53 |
| Number of frames per exposure | 40 |
| Energy filter slit width (eV) | 20 |
| Data collection software | EPU |
| Number of exposures per hole | 2 |
| Defocus range (μm) | -1.5 to -2.5 |
| Number of micrographs collected | 4024 |
| Number of micrographs used | 3923 |
| Number of initial particles | 3,388,883 |
| Symmetry | C1 |
| Number of final particles | 139,718 |
| Resolution (0.143 gold standard, Å) | 3.34 |
| **Atomic model refinement** | |
| Software | phenix |
| Clashscore, all atoms | 22.36 |
| Poor rotamers (%) | 0 |
| Favored rotamers (%) | 94.14 |
| Ramachandran outliers (%) | 0 |
| Ramachandran favored (%) | 85.34 |
| MolProbity score | 2.5 |
| Bad bonds (%) | 0 |
| Bad angles (%) | 0.02 |
| RNA Bad bonds (%) | 0 |
| RNA Bad angles (%) | 0 |

**Extended Data Table 3. Cryo-EM data collection, processing, and model validation of the NSUN2-pre-tRNA$^{Leu}$-SAM complex.**

| | NSUN2-pre-tRNA$^{Leu}$-SAM complex |
|---|---|
| **Data collection and processing** | |
| Microscope | Titan Krios G4 |
| Voltage (kV) | 300 |
| Camera | Falcon4i |
| Grid type | R 2/1 Quantifoil copper grid (200 mesh) |
| Magnification | 130,000× |
| Pixel size (Å) | 0.97 |
| Total exposure ($e^-$/Å$^2$) | 50 |
| Exposure time (s) | 4.19 |
| Number of frames per exposure | 40 |
| Energy filter slit width (eV) | 20 |
| Data collection software | EPU |
| Number of exposures per hole | 4 |
| Defocus range (μm) | -1.5 to -2.5 |
| Number of micrographs collected | 4250 |
| Number of micrographs used | 4100 |
| Number of initial particles | 3,390,207 |
| Symmetry | C1 |
| Number of final particles | 112,021 |
| Resolution (0.143 gold standard, Å) | 3.30 |
| **Atomic model refinement** | |
| Software | phenix |
| Clashscore, all atoms | 23.41 |
| Poor rotamers (%) | 0 |
| Favored rotamers (%) | 75.56 |
| Ramachandran outliers (%) | 0 |
| Ramachandran favored (%) | 97.38 |
| MolProbity score | 2.46 |
| Bad bonds (%) | 0 |
| Bad angles (%) | 0.02 |
| RNA Bad bonds (%) | 0 |
| RNA Bad angles (%) | 0 |

**Extended Data Table 4. Cryo-EM data collection and processing of the NSUN2- tRNA$^{Lys}$-SAM.**

| | NSUN2- tRNA$^{Lys}$-SAM complex |
|---|---|
| **Data collection and processing** | |
| Microscope | Titan Krios G4 |
| Voltage (kV) | 300 |
| Camera | Falcon4i |
| Grid type | R 1.2/1.3 Quantifoil copper grid (200 mesh) |
| Magnification | 130,000× |
| Pixel size (Å) | 0.93 |
| Total exposure ($e^-/Å^2$) | 50 |
| Exposure time (s) | 5.82 |
| Number of frames per exposure | 40 |
| Energy filter slit width (eV) | 20 |
| Data collection software | EPU |
| Number of exposures per hole | 2 |
| Defocus range (μm) | -1.5 to -2.5 |
| Number of micrographs collected | 5548 |
| Number of micrographs used | 5427 |
| Number of initial particles | 2,716,257 |
| Symmetry | C1 |
| Number of final particles | 67546 |
| Resolution (0.143 gold standard, Å) | 3.93 |
| **Atomic model refinement** | |
| Software | phenix |
| Clashscore, all atoms | 34.41 |
| Poor rotamers (%) | 0 |
| Favored rotamers (%) | 89.90 |
| Ramachandran outliers (%) | 0 |
| Ramachandran favored (%) | 87.61 |
| MolProbity score | 2.73 |
| Bad bonds (%) | 0 |
| Bad angles (%) | 0 |
| RNA Bad bonds (%) | 0 |
| RNA Bad angles (%) | 0 |
